# Topological Gelation of Reconnecting Polymers

**DOI:** 10.1101/2022.09.21.508941

**Authors:** Andrea Bonato, Davide Marenduzzo, Davide Michieletto, Enzo Orlandini

## Abstract

DNA recombination is a ubiquitous process that ensures genetic diversity. Contrary to textbook pictures, DNA recombination, as well as generic DNA translocations, occur in a confined and highly entangled environment. Inspired by this observation, here we investigate a solution of semiflexible polymer rings undergoing generic cutting and reconnection operations under spherical confinement. Our setup may be realised using engineered DNA in presence of recombinase proteins or by considering micelle-like components able to form living (or reversibly breakable) polymer rings. We find that in such systems there is a topological gelation transition, which can be triggered by increasing either the stiffness or concentration of the rings. Flexible or dilute polymers break into an ensemble of short, unlinked and segregated rings, whereas sufficiently stiff or dense polymers self-assemble into a network of long, linked and mixed loops, many of which are knotted. We predict the two phases should behave qualitatively differently in elution experiments monitoring the escape dynamics from a permeabilised container. Besides shedding some light on the biophysics and topology of genomes undergoing DNA reconnection *in vivo*, our findings could be leveraged *in vitro* to design polymeric complex fluids, e.g., DNA-based complex fluids or living polymer networks, with desired topologies.

---

Recombination of genetic material involves the transient cleavage of two DNA segments that are spatially proximate in 3D – although not necessarily adjacent in 1D – followed by alternative rejoining of DNA ends. Beyond its role in meiosis [1], similar topological processes involving the reconnection of DNA segments are also seen in the proliferation of transposable elements [2, 3] and the integration of viral DNA in the host genome [4]. More recently, artificially driven DNA translocation- and recombination-like events have been used to map highly-accessible genomic sites [5] and scramble synthetic yeast chromosomes [6]. Recombination operations on a plasmid *in vitro* are known to yield linked or knotted DNA products [7–9]. This observation suggests that unrestricted DNA recombination *in vivo* may pose a pressing topological problem to the cell [10, 11], but also that recombination may be employed to design topologically non-trivial DNA molecules.

Enzyme-mediated recombination has been well studied on short plasmids in dilute conditions [8, 12–15]. On the other hand, the topological consequences of recombination operations on long and entangled DNA are far less investigated or understood. Inspired by this problem, here we study a system of ring polymers continuously undergoing cutting and reconnection operations – hereafter called “reconnecting” or “recombinant” rings – inside a sphere (Fig. 1A). We note that, at variance with meiotic recombination, where two finite chromosome sections are exchanged, our reconnection operations are performed by introducing an exchange event on a single site, followed by alternative rejoining of the polymer segments (see Fig. 1A). Therefore, our model entails a highly simplified view of recombination, and its aim is consequently limited to exploring the generic and qualitative topological feature of recombination in confinement, rather than making quantitative predictions. At the same time, our system can be viewed more generally as a confined melt of living polymer loops. Living polymers are reversibly breakable: like the polymers in Fig. 1A they can break and rejoin locally (i.e., reconnect) whilst remaining in thermal equilibrium [16]. A melt of living polymer loops can form spontaneously following polymerisation of monomers, given an appropriate choice of the reaction kinetics [17]. As we discuss below, such melts can in principle be realised experimentally, hence are potentially relevant to materials science.

**FIG. 1.**
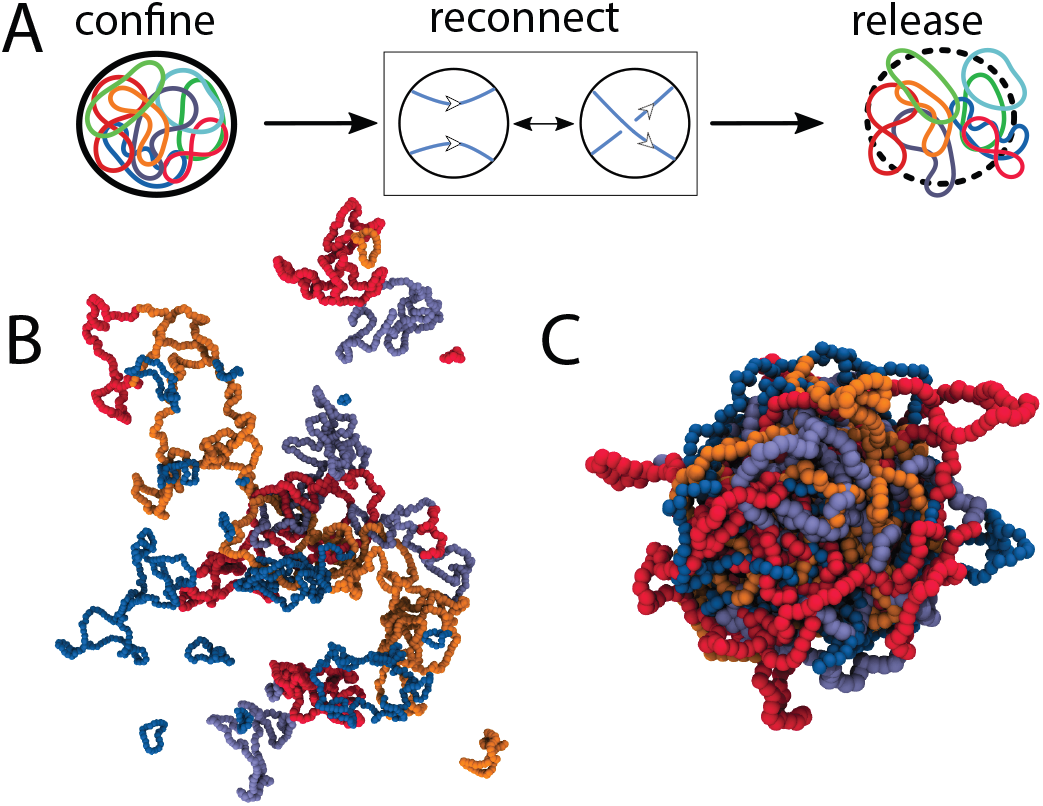
Phases of reconnecting rings. **A** We study ring polymers allowed to recombine/reconnect within a sphere of radius *R*. **B-C** The two panels show two possible states of the system at equilibrium and after relaxing the confinement. **B** Sketch of an ensemble of many, small and mostly unlinked rings whereas **C** shows an ensemble of few, long and linked rings. The snapshots are taken after releasing confinement for ease of visualisation.

We discover that, depending on polymer stiffness and radius of the confining sphere, these systems display a topological transition between a regime with many short, unlinked and segregated rings and another one with few long, mixed and linked rings. Geometrically, this transition is naturally explained as the result of a competition between bending energy of the loops and entropy of the system. Topologically, it can be seen as a gelation transition and understood in terms of the critical overlap concentration *c*^*\**^, above which linking is expected to be entropically favoured. Our gel of recombinant polymers is fundamentally different from other types of topological gels formed by nearly monodisperse loops in the presence of topoisomerase-like enzymes, such as the kinetoplast DNA network [18] or Olympic gels [19, 20], because in our case multicomponent links are generically polydisperse and typically contain one or very few loops which are much longer than the rest, and are often knotted. Similarly, our setup is distinct from that of previous works investigating the segregation of fixed-size polymer rings in a melt or under confinement [21–28], because the latter did not consider reconnection operations, which can change loop sizes and global topology.

The topological gelation we find in this work suggests that unregulated single-site reconnection of DNA *in vivo* should be highly detrimental. On the other hand, being able to construct a phase diagram for gelation *in vitro* can be useful from a materials science perspective, as it may provide an avenue to design experiments with DNA rings which undergo recombination under confinement to yield linked and knotted products with desired topologies, or materials with topologically controlled mesoscopic properties, e.g. Olympic ring-like gels [19, 20].

## A MODEL FOR RECONNECTING POLYMERS

Unless otherwise specified, the system is initialised as one ring with *N* = 1000 beads of size *σ* (see SI for results with *N* = 10000). The beads interact via a purely repulsive Lennard-Jones potential, and adjacent beads are connected by FENE bonds. A key parameter of our system is the stiffness *K* of the chains, which is proportional to the chain persistence length, imposed via a Kratky-Porod potential [29, 30] (see Methods). The simulations are performed in LAMMPS [31] using a Langevin thermostat and a timestep Δ*t* = 0.001*τ*_*B*_, with *τ*_*B*_ the Brownian time (see SI for more details). The polymer is initialised inside a large sphere that is slowly compressed to *R* = 7*σ* and subsequently allowed to equilibrate. After this equilibration step, we allow the ring to undergo reconnections, i.e., transient single-site breakage followed by alternative joining, between any two segments that are proximal in 3D (Fig. 1; effectively we consider only segments closer than *r*_*c*_ = 1.3*σ*). Reconnection moves are attempted at every integration step (if the distance condition is satisfied) and are performed via a modified `fix bond/swap` [32–35] (see SI). Since we accept or reject the moves according to a Metropolis test, the actual reconnection rates *κ*_*r*_ depend on the stiffness parameter *K*: for instance, 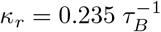 at *K* = 0 and 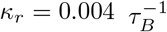 at *K* = 5. By mapping our Brownian time to real units (assuming *σ* = 30 nm and a medium with viscosity *η* = 100 cP as the nucleoplasm) these rates can be converted to 0.5 − 30 s^*−*1^. These are overall faster than the 1*/*min recombination rates in biological systems [36], but of the same order as recombination in wormlike micelles [16, 37]. Importantly, the precise value of *κ*_*r*_ does not affect our conclusions as we are interested in the longtime, steady state behaviour of the system, rather than the transient dynamics.

Every 10^2^*τ*_*B*_ = 10^5^Δ*t* we take a snapshot of the system and reconstruct its topology, record the chain number, *N*_*r*_(*t*), as well as their length distribution *L*_*r*_(*n, t*). The total number of beads in the system is kept constant to 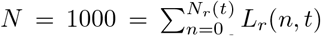, so that the monomer density is *ρ* = 3*N/*(4*πR*^3^) ≃ 0.7*σ*^*−*3^, or equivalently, the volume fraction *ϕ* = 0.37. We have further checked that our system yields similar results with *N* = 10000 beads and also starting from different initial conditions. For more details on our model, see Materials and Methods and SI.

## RESULTS

### Geometry of Confined Reconnecting Rings

We begin by characterising the geometrical features of the system as a function of the stiffness parameter *K* (proportional to the persistence length of the rings, *l*_*p*_), for a fixed value of spherical confinement *R*. We monitor the average number of rings and their average length: in our simulations, both these quantities evolve to reach a well-defined steady state value (see Fig. 2A,B).

**FIG. 2.**
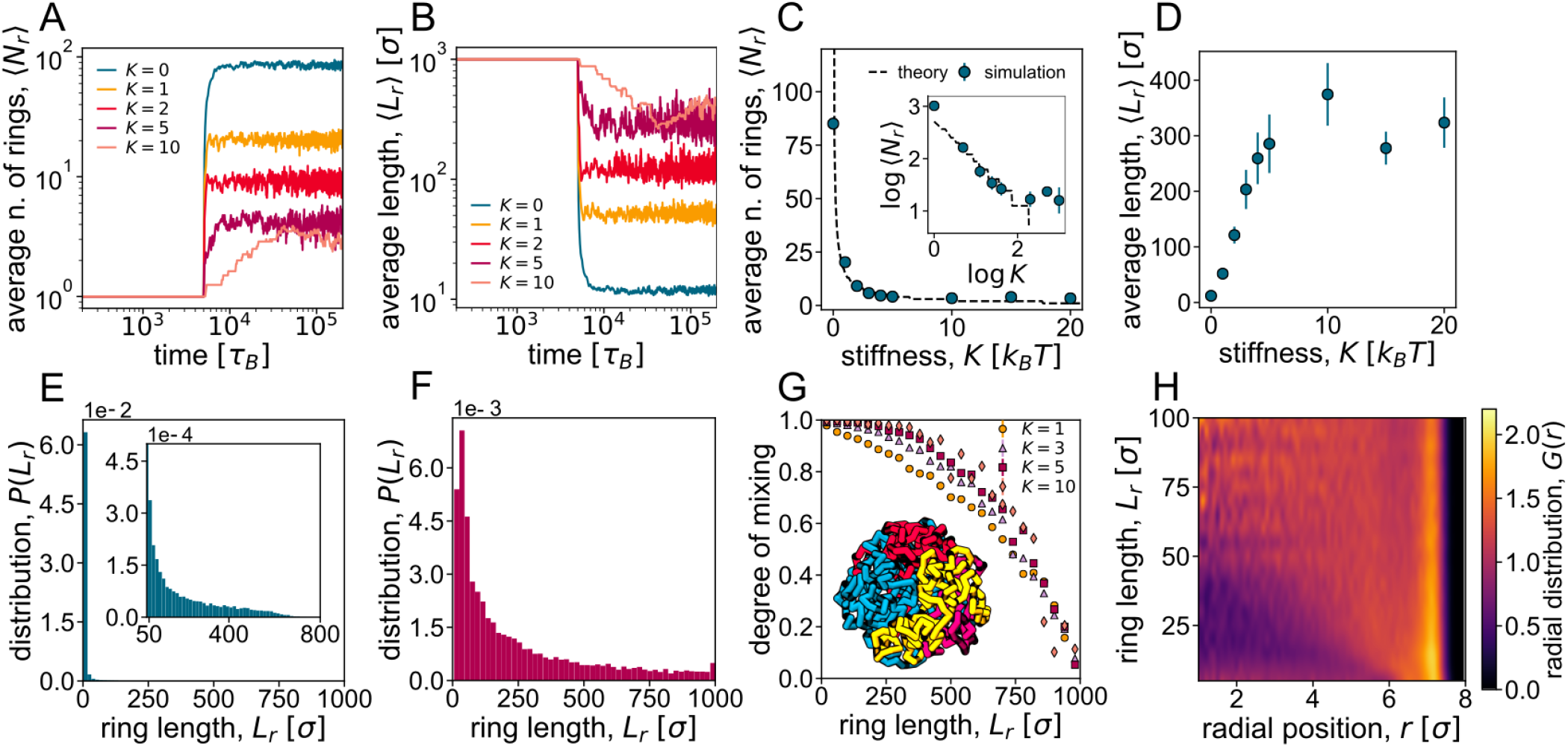
Geometrical transition of reconnecting rings. **A-B**. Time evolution of the average number of reconnecting rings *N*_*r*_ **(A)** and their average contour length *L*_*r*_ **(B)**. Different curves refer to different values of the stiffness parameter *K*. All the data refer to a spherical confinement of radius *R* = 7*σ*. **C-D**. Average steady-state values of ⟨*N*_*r*_⟩ and ⟨*L*_*r*_⟩, respectively, as a function of *K*. The black line in panel **(C)** and inset shows the theoretical prediction from (1) with *λ* = 5.3 and *a* = 0.27. **E.-F**. Distributions of ring size in steady state, for *K* = 1 **(E)** and *K* = 5 **(F)**, indicating that the system is always highly polydisperse. **G**. Probability that a ring of length *L*_*r*_ mixes with other rings as a function of *L*_*r*_, for different values of *K*. **H**. Heatmap of the radial distribution (*G*(*r*)) of monomers in rings of length *L*_*r*_ as a function of *r*, the distance from the centre of the confinement sphere, and *L*_*r*_.

Our data are suggestive of a smooth transition or crossover between a regime in which many short rings populate the sphere (at low stiffness *K*) and another one in which few long rings remain in steady state (at high stiffness *K*). We temporarily refer to these as the short ring and the long ring regimes respectively. A typical snapshot of the system in the two regimes is shown in Fig. 1B-C (where the confining sphere is removed for ease of visualisation). The transition can be understood in terms of the competition between the bending energy of the loops – regulated by *K*, which favours the long ring regime – and their combinatorial and translational entropy – which favours the short ring regime. To estimate entropy, we note that if we allow all rings to freely reconnect, the number of ways in which *N* beads can be distributed into *m* rings (without leaving any one of these empty) is given by the Stirling number of the second kind

{*N, m*} ∼ *m*^*N*^ */m*! for large *N*. Including the bending energy as well as the configurational entropy of rings (see SI for details), the total free energy of the system – apart from an irrelevant additive constant – can be estimated as

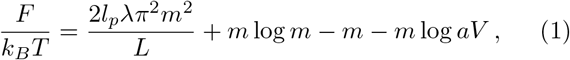

with *λ* a numerical factor related to the specific shape taken by a curved polymer and *a* a numerical factor specified in the SI. We note that we assumed the curvature to scale as ∼ 1*/L*, which is valid for short rings such that *L* ≃ *ml*_*p*_; we shall see below that this assumption holds for our system.

By minimising the free energy with respect to *m*, we can find the average number of rings as a function of *K* and *R*. As seen in Fig. 2C (and inset), this mean field theory captures the numerical results for the average number of rings, ⟨*N*_*r*_⟩, very accurately. In particular, both our simulations and our theory predict that ⟨*N*_*r*_⟩ ∼ *K*^*−*1^. (1) also naturally describes the behaviour of the expected mean length ⟨*L*_*r*_⟩, thanks to the fact that the total number of monomers is conserved. As shown in Fig. 2D, the mean length displays a smooth transition, reaching a plateau at large values of *K*. Therefore our theory shows that increasing *K* leads to a transition (or crossover) between the short ring and the long ring regimes. In the SI we give more details about our semi-analytical theory, and show that a similar transition can be observed at fixed stiffness *K* by decreasing the sphere radius *R* (Fig. S1).

An intriguing feature of our reconnecting polymer system is that the rings obtained in steady state have a very broad size distributions (see Figs. 2E,F and Figs. S2,S3). Notably, this is true for both small and large *K*, and the size distribution is a power law, *P* (*L*_*r*_) ∼ 1*/L*_*r*_, for all cases (Fig. S2). As reconnection changes polymer length but does not violate detailed balance in our model, the system is effectively in thermodynamic equilibrium, and the size distribution should be linked to the Boltzmann weight of rings of different sizes. Neglecting the dependence on *K*, which is expected to be a fair approximation for sufficiently long rings, we therefore expect the size distribution to be 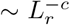, i.e., the probability of forming a loop of size *L*_*r*_. We note that a power-law size distribution of rings is also found, for analogous reasons, in living (reversibly breakable) polymers in the phase where loops are favoured over linear chains [17]. As discussed in more detailed later, the inherent polydispersity of reconnecting rings plays an important role to determine the emerging macroscopic properties of our system.

### Radial Positioning and Mixing of Reconnecting Rings

*In vivo*, chromosomes segregate into territories which position themselves non-randomly with respect to the nuclear lamina. In this process, entropic effects play an important role [38, 39]. Motivated by this, we ask how entropy and stiffness affect mixing and non-random positioning of reconnecting rings (which may be viewed as toy chromosomes) inside the sphere (a toy nucleus).

To this end, we construct a parameter, similar to the conditional entropy used to measure the entropy of mixing [40, 41], which quantifies the probability of finding monomers from other rings within a sphere centred at a monomer in a given ring. In Fig. 2G, we show that this mixing probability decreases with ring length and increases with stiffness *K*, as expected for concentrated solutions [42]. In other words, longer and more flexible rings are less mixed. To further characterise the spatial arrangement of the rings, we compute the normalised radial density of monomers, *G*(*r*) (Figs. 2H and S4). Uniform positioning of beads, and hence of rings, within the sphere corresponds to a constant *G*(*r*). Instead, we see that smaller loops (small *L*_*r*_) are depleted in the interior of the sphere and enriched at the periphery *r* ≃ 7*σ* (Figs. 2H and S5). This is a consequence of steric depletion [43] and geometry, as smaller soft objects can approach a surface more frequently than larger ones. This finding also suggests that smaller plasmids or extra-chromosomal DNA may be found more frequently towards the periphery of the cell (in bacteria) or nucleus (in eukaryotes).

### The Short and Long Ring Phases are Separated by a Topological Gelation Transition

Up to now, we have shown that increasing the stiffness *K* in a melt of reconnecting rings drives a transition or crossover between a short ring and a long ring regime. As anticipated, and as shown in the SI, the same transition or crossover can also be achieved by increasing the ring density at a fixed value of the stiffness. More insight into this transition, and its underlying physical mechanism, can be gained by analysing the overlap between reconnecting rings of different stiffness. For increasing values of stiffness, the average length of the rings in steady state is larger (see Fig. 3A, blue circles are the same as in Fig. 2D) in turn entailing more overlaps. Calling *L* the average length of the rings, we can define a critical overlap concentration as

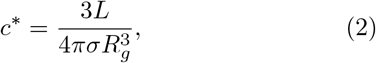

with *R*_*g*_ the radius of gyration of the rings. This is the concentration above which rings of size *L* start to overlap with each other. For our case, assuming that 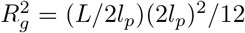 for *K* ≥ 1, and that 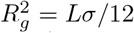 for *K* = 0, we find *c*^*\**^ as a function of *L* (Fig. 3B). Using this curve, we can predict the critical value of the length for which *c* = *c*^*\**^ as a function of stiffness, in turn yielding the red curve in Fig. 3A. This curve is a decreasing function of *K*: in other words, increasing *K* makes it easier for rings to feel and overlap with each other.

Importantly, as a consequence of the opposing dependencies of the average ring length and of the critical overlapping length as a function of *K*, there is a point at which the two curves cross (Fig. 3A). The crossing point marks the critical point separating the short ring from the long ring regime. In more detail, when the red curve (filled squares) in Fig. 3A is above the blue curve (filled circles), then *c < c*^*\**^, and we expect rings should not overlap and therefore segregate, leading to the short ring phase (left panel of Fig. 1B). Instead, when the red curve is below the blue one, then *c > c*^*\**^, and we expect rings to mix and reconnect significantly with each other, leading to the long ring phase (right panel of Fig. 1B). This reasoning suggests that the transition we observe is akin to a gelation transition, where rings can be seen as soft particles which start to interact once *c/c*^*\**^ becomes large enough. It is indeed natural to expect that the system should behave as a gas or liquid of soft particles (the rings) for *c < c*^*\**^, and as a gel (or a solid-like structure) for *c* sufficiently larger than *c*^*\**^. The main difference with respect to colloidal gels of soft particles is that in our case there are no direct attractive interactions between rings. However we expect topological interactions to be present, as above *c*^*\**^ rings can link with each other, as occurs in concentrated solutions of fixed-size crossable rings [44]. For this reason we refer to the transition between the short and long ring phases as *topological gelation*, and we shall reinforce this interpretation in the analysis described in the following sections. We note that in our system increasing *K* leads to a decrease in *c*^*\**^, whereas a decrease in *R* leads to an increase in *c*. This is why the system can be made to gel either by increasing *K* (see Fig. 3), or by decreasing *R*, because both these variations increase the value of *c/c*^*\**^, hence favour the gel phase.

**FIG. 3.**
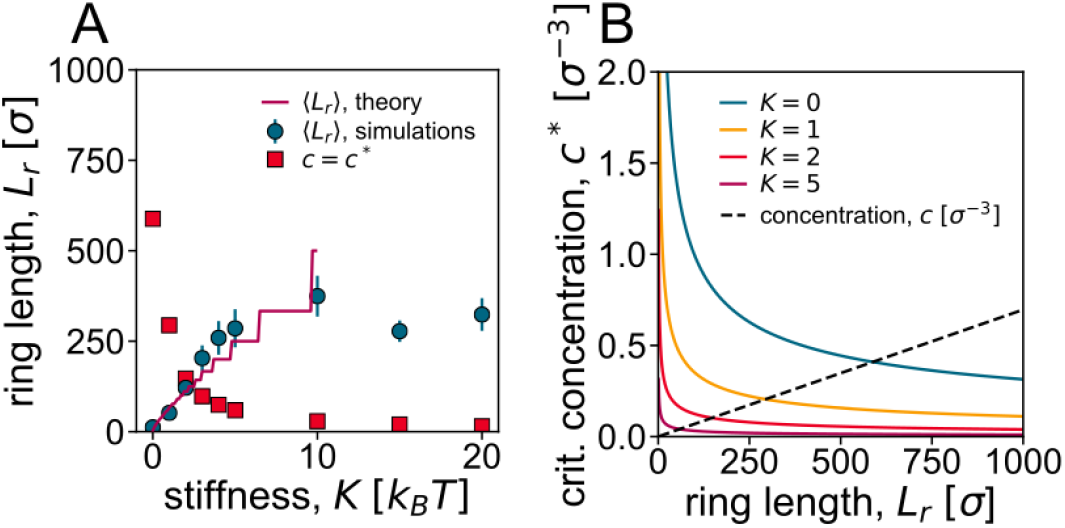
Critical point for gelation. **A**. Filled squares: critical length *L*^*\**^, for which the concentration *c* of a ring in a sphere with *R* = 7*σ* equals *c*^*\**^, as a function of *K*. Filled circles: mean ring length versus *K*. The critical point for gelation is expected to be where the two curves cross. **B**. Calculation of *L*^*\**^. At *L*^*\**^, the curves (*c*^*\**^ as a function of *K*) and the dotted line (*c* as a function of *K*) intersect. *c*^*\**^ is predicted for an ideal polymer ring (2).

We note that, for *K* = 5, an isotropic-to-nematic transition is expected around monomer density *ρ* = 0.85*σ*^*−*3^ [29, 30], which is larger than the monomer density at which we work at *ρ* = 0.7*σ*^*−*3^. While alignment effects may be important, we argue that the main role in driving the topological transition is played by the overlap concentration *c*^*\**^, as explained above. In line with this, in the SI we show that a similar unlinked-to-linked transition is observed for fully flexible (*K* = 1) chains, albeit at larger densities *ρ* ≃ 0.76*σ*^*−*3^.

### Topological Gelation is Accompanied by the Formation of a Percolating Network of Linked Loops

We now discuss in more depth the topological nature of the gelation transition between the short and long ring phases. To quantify the topological entanglement between the rings in the system, we first compute the Gauss linking number (see Materials and Methods). We find that, as the reconnecting rings get stiffer, the typical topologies found at fixed confinement radius *R* are markedly different. Figs. 4A,B show the time dependence of the number of linked pairs and of ⟨|*Lk*|⟩, the total absolute value of the linking number (see Materials and Methods and SI for its precise definition). These curves, and the steady-state averages plotted in Figs. 4C,D, show that both these quantities increase with the stiffness *K*, as expected from our argument that the short-to-long ring transition is akin to gelation (Fig. 3). Flexible rings are therefore typically short and unlinked, whereas stiffer loops are typically longer and linked (see Fig. S6 for the ⟨|*Lk*|⟩ distribution). The transition or crossover which we observe by increasing stiffness is thus associated with an increase in topological entanglement and in linking between chains. In keeping with our interpretation of the long-ring phase as a gel, we expect that the topological entanglement between rings should endow our system with a non-zero elastic modulus.

**FIG. 4.**
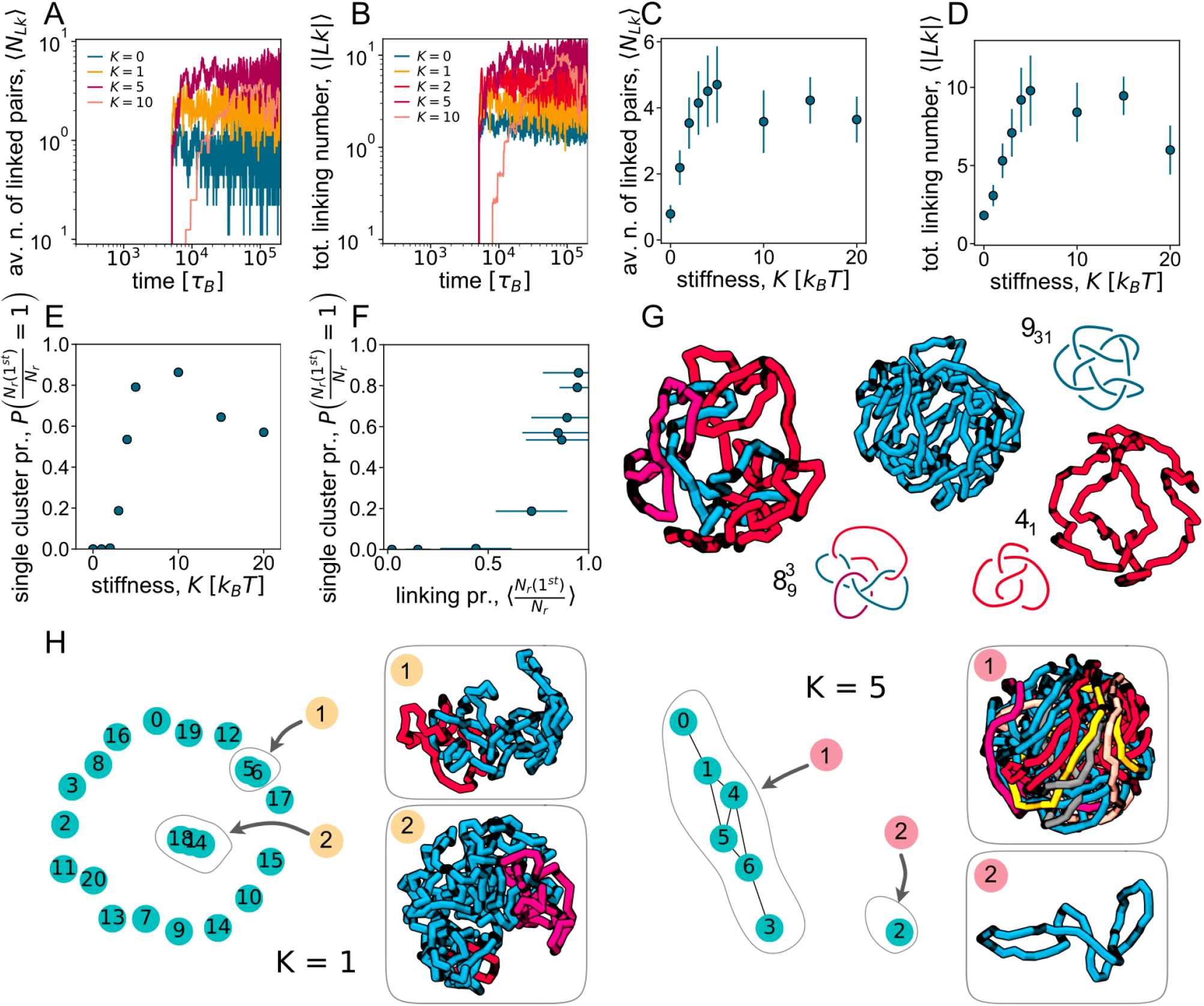
Topological gelation. **A-B**. Time dependence of the number of linked pairs of rings *N*_*Lk*_ **(A)** and average of the total unsigned linking number, ⟨|*Lk*|⟩, **(B)**. Different curves refer to different values of the stiffness parameter *K*. **C-D**. Average steady-state values of *N*_*Lk*_ and ⟨|*Lk*|⟩ respectively as a function of *K*. **E-F**. Probability of observing a single cluster as a function of *K* **(E)** and as a function of the linking probability **(F). G**. Examples of knots and catenanes found in the steady-state configurations at *K* = 5. **H**. Snapshots of clusters of linked rings in simulations with *K* = 1 and *K* = 5. Each sampled configuration is associated with a network; linked rings (vertices) are connected by edges. A connected component of the network represents a cluster of linked rings.

As topological gelation is approached, the nature of the typical network of linked rings forming in steady state changes qualitatively, as shown in Figs. 4E-F. For low *K* (in the liquid phase), the network has low connectivity, and clusters have typically one or very few nodes (Fig. 4E). In sharp contrast, for large *K* (in the gel phase), there is a large connected component which accounts for a substantial fraction of all rings (Fig. 4F). More precisely, we can quantify topological gelation by measuring the probability that the network condenses into a single connected cluster of nodes, which appears to depart from 0 abruptly at a critical value of *K* (Fig. 4F). From Figs. 4E-F, one can appreciate a dramatic stepwise change at the critical value *K* ≃ 2, which is where *c/c*^*\**^ ≃ 1 according to our theory (see Fig. 3A). Gelation is therefore accompanied by a percolation transition in the network of linked rings. In our system, rings outside the largest connected component are typically unlinked, so that the average fraction of rings in the largest component is approximately equal to the linking probability for any ring in the network. Fig. 4F shows that gelation occurs when this linking probability proxy is about 1*/*2; interestingly, this is equal to the bond percolation threshold for the square lattice, but significantly higher than that of most standard 3D lattices.

### Rings in Reconnecting Topological Gels often Form Complex Knots

We now turn to a more detailed analysis of the topology of the gel phase which, as we shall see, features some unique properties that cannot be found in topological gels with strand-crossing rings of fixed size. While classic Olympic gels [19, 20] are typically made by singly-linked monodisperse rings (see also kinetoplast DNA [18, 44, 45]), topological gels from confined reconnecting rings possess much more complex and exotic structures (Fig. 4G-H). First, recombination of rings can create knots and, accordingly, we find that often some rings in our gel phase are knotted. We typically observe that only the longest component in a link is knotted. Such knots can be relatively complicated – for instance one of the two examples featured in Fig. 4G can be identified as a 9_31_ knot. Second, even unknotted rings may form complex catenanes: the one shown in Fig. 4G displays three rings that form an 8-crossing link (note only two of the rings are pairwise linked, as a Solomon knot). Results for larger systems (where longer chains are confined in a larger sphere, Fig. S8) show even more exotic. For instance, Fig. S8 shows a circular catenane, and a 7_5_ knot linked to two unknots in two different ways – as a Hopf link with one, and as a Solomon knot with the other. As found for the total linking number ⟨|*Lk*|⟩, we also observed that the complexity in topology tends to increase with *K*, or *c/c*^*\**^, so that deeper in the gel phase topologies are more complex than close to gelation.

How do such complex topologies form spontaneously through recombination/reconnection? We argue that this is due to the fact that the reconnection process allows ring length to vary. Indeed, in dilute conditions, multicomponent links made of equal-size rings which can exchange material are unstable for entropic reasons, and as a result one of the rings expands at the expense of all the others [46]. Our results suggest that a similar entropic drive leads to the growth of a single ring in a multi-component link also under confinement. Then, the larger ring is more likely to contain a knot, while the shorter rings are more likely to be unknotted; this is because knots have a statistical size which increases with their topological complexity [47]. Clearly, this argument shows why topologies like those in Figs. 4H and S8 are impossible to obtain starting with monodisperse rings subject to strand-crossing moves, as these do not allow material exchange between rings.

### Topological Gelation Traps the Reconnecting Rings inside a Permeabilised Sphere

The topological transition we uncovered in the previous section can be vividly appreciated by the following procedure, which is inspired by elution experiments in biophysics and could in principle be realised in the lab [48]. After a steady state is reached in our simulations, we permeabilise the confining sphere by converting it into a spherical mesh with pores of controlled size. We then disallow further reconnection and monitor the number of monomers still inside the sphere as a function of time, *n*(*t*). The curves *n*(*t*) are reported in Fig. 5A,B for the liquid (*K* = 0) and gel (*K* = 3) regimes, respectively. It is apparent from the markedly different curves that the two regimes lead to very different escape dynamics. Note that, to single out the effect of the different topological states, after permeabilisation we set the stiffness to the same value (*K* = 1) in all cases.

**FIG. 5.**
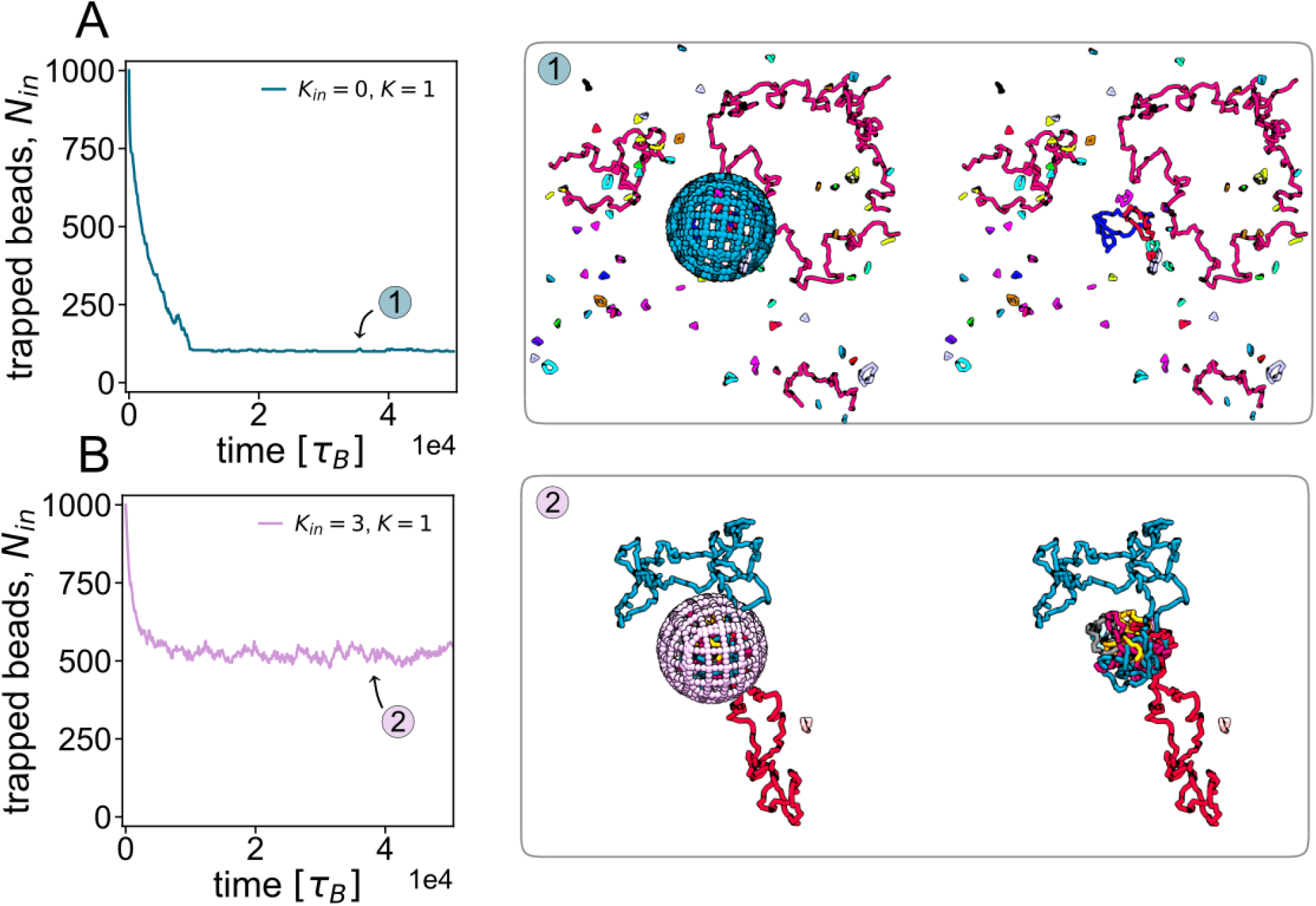
Escape dynamics of reconnecting rings from a permeabilised sphere. Results of simulated elution experiments from a permeabilised sphere. The different structures correspond to a system of reconnecting rings with different *K*. After permeabilising the sphere, *K* was set to 1 for all systems and the reconnection was disallowed, in order to focus on the effect of topology on the escape dynamics. Left column: number of monomers inside the sphere as a function of time for an initial value of *K* equal to 0 **(A)** and 3 **(B)**. Corresponding snapshots are shown on the right (with and without the sphere to ease visualisation of the topological structures). Simulations are performed inside a box with periodic boundaries.

In the liquid phase, the “dust” of overwhelmingly small and unlinked rings rapidly diffuses out of the sphere, as their translational entropy increases if confinement is removed, and such rings are small enough to translocate through the pores (see snapshots in Fig. 5A). In this liquid or gas phase, longer rings, if present, are typically unlinked and can eventually exit the sphere. This behaviour is qualitatively equivalent to that observed in experiments in permeabilised cells which show elution of sufficiently small molecules, such as diffusing proteins or DNA fragments of small size [48].

In sharp contrast, when the system is in the gel phase, the network of linked loops which emerges after reconnection is too large and topologically complex to translocate through the pores; most of the system is thus kinetically trapped inside the permealised sphere. This is apparent from the plateau of *n*(*t*), suggesting a long-lived steady state with a substantial proportion of rings still inside the sphere (see Fig. 5B). This second scenario is also reminiscent of elution experiments, where large superstructures, like microphase separated protein droplets or chromatin-protein aggregates such as transcription factories, resist elution and remain inside the permeabilised nuclei [48].

In the SI, we also consider the case where reconnections are still possible, and the stiffness is not reset, after permeabilisation (Fig. S9). This situation may correspond more closely to an actual experiment with confined living polymer rings. While the escape dynamics is again vastly different in the liquid and gel phase, in this case all rings escape the sphere at small *K*, and the number inside the sphere decays as a stretched exponential. Instead, the behaviour in the gel phase is very similar to that shown in Fig. 5, and the value of *n*(*t*) reaches a non-zero plateau at large times.

## DISCUSSION AND CONCLUSION

In summary, here we have used coarse-grained molecular dynamics simulations to study the behaviour of a solution of polymer rings undergoing recombination-like reactions (which we call reconnections, Fig. 1) inside a spherical container. There are two potential avenues to realise this system experimentally and to test our theoretical predictions *in vitro*. First, the system may be recreated by using a concentrated or confined mixture of suitably engineered DNA plasmids and recombinase enzymes. We note that a similar system was built to create synthetic scrambled yeast chromosomes [6]. Second, one may use a confined ensemble of living ring polymers. The latter may be realised, for instance, by using fibril-forming proteins or patchy particles above the critical micellar concentration. As in wormlike micelles rings are normally absent or irrelevant [16], it would be necessary to select proteins or particle whose geometry favours the formation of loops [49, 50]. Additionally, the system we have considered here may be used as a very simplified framework to model the behaviour of recombinant DNA *in vivo* (e.g., in the scrambled yeast chromosome system [6]).

Our main finding is that a confined solution of reconnecting polymer rings harbours a transition, or crossover, between two fundamentally distinct regimes, which can be triggered either by stiffening the polymers, or by increasing their density. For flexible or sufficiently dilute polymer rings, reconnection results in the production of a gas, or fluid, or short segregated and unlinked loops. For polymers with a sufficiently large persistence length, or for a sufficiently dense solution, a network of long linked loops emerges in steady state. This transition is a topological analogue of the gelation transition observed for sufficiently dense suspensions of sticky colloids, where the formation of force chains is substituted by topological linking. Like gelation and vitrification, our topological transition is accompanied by a dramatic slowdown in the system dynamics, which could be quantified experimentally, for instance, by measuring the rate of escape from the confining sphere when the latter is pierced by appropriate-size pores to permeabilise it, as in elution experiments with DNA or chromatin. For large enough stiffness, our system is expected to undergo an isotropic-to-nematic transition involving spooling [51]. Arguably, nematic alignment of polymer segments in this system may prevent proficient linking between different loops. Indeed, our simulations suggest that the onset of the topological gelation is mainly determined by the overlap concentration *c*^*\**^, which depends on the equilibrium size of the polymer loops (itself dependent on the stiffness). Accordingly, in SI we show that the unlinked-linked transition can be seen also for very flexible chains (*K* = 1) at large enough volume fractions.

The recombinant topological gel we have found here is fundamentally distinct from other topological gels obtained in DNA ring solutions either *in vitro* or *in vivo* by using topoisomerase, such as Olympic gels [19, 20] or the kinetoplast DNA [18, 44, 45]. In these two cases, DNA rings are monodisperse in length, typically unknotted and singly linked. In our recombinant gel, instead, rings are polydisperse, with a broad length distribution (see Fig. S2); they are also often knotted (see Fig. S10), especially at high concentrations, or under tight confinement. The difference between these two types of topological gels is inherently due to the difference between recombinase-like and topoisomerase-like operations, as the former does not need to preserve the ring lengths.

Besides being of fundamental interest as an example of a topological phase transition in a soft condensed matter system, our results can therefore be exploited to design DNA gels with complex topologies. In this respect it is important to note that topological (knot or link) complexity tends to increase with stiffness, or confinement, so that by selecting parameters appropriately it should be possible to sieve networks with desired topologies. We anticipate that topological gelation can also be found if confinement is replaced by crowding, for instance by varying the stiffness of a suspension of recombinant polymer loops of a sufficiently high volume fraction. In such a geometry, the transition could be characterised, for instance, by measuring the bulk rheology response of the system, as the elastic modulus should be non-zero in the gel phase.

In the context of recombinant DNA *in vivo*, we speculate our results suggest that recombination, when left unchecked, is likely to create a topological gel given the high genomic density found under physiological conditions in the nuclei of living cells. Gelation is likely detrimental for the cell, as it would lead to entanglements hindering, for instance, chromosome segregation during cell division. Interestingly, specific biological mechanisms, such as Topoisomerase action and active chromosome movement, are indeed in place to remove chromosome interlocks [52–54], which are recombination-driven entanglements which form during early meiosis [55].

We hope that our work will inspire and inform future experiments on reconnecting DNA plasmids or living polymer rings at large density.

## I. METHODS

We simulate confined bead-spring polymers made of beads of size *σ* and mass *m* at a temperature *T*, inside a spherical container of radius *R*. The equations of motion, force fields, and the implementation of the reconnection moves are described in the SI.

An attempted reconnection move which would change the configuration of the system from *ω* to *ω*^*′*^ (see Fig. 1A and Model section) is accepted with probability

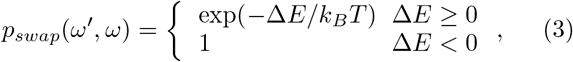

where Δ*E* is the energy difference between *ω* and *ω*^*′*^.

The pairwise topological complexity of the system of rings is estimated by computing the Gaussian linking number for each pair of rings *γ*_*i*_ and *γ*_*j*_, which is given by

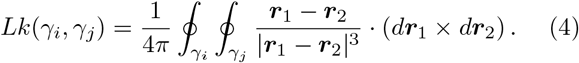

For each pair (*i, j*) and sampling time *t* we defined

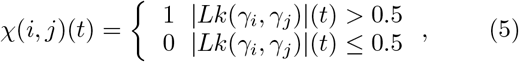

and, by summing over all pairs, we obtained the number of linked pairs of a given configuration at time t:

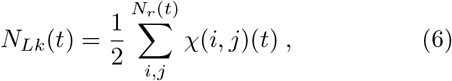

where *N*_*r*_(*t*) is the number of rings at time *t*. The total absolute value of the linking number of a configuration is computed as

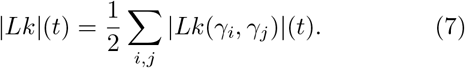

Different averages of *N*_*Lk*_ and |*Lk*| are shown in Figs. 4A-D. Additional model details and supporting results are given in the SI.

## II. ACKNOWLEDGEMENTS

DMi acknowledges the ERC and Royal Society for funding. This project has received funding from the European Research Council (ERC) under the European Union’s Horizon 2020 research and innovation programme (grant agreement No 947918, TAP).

